# Pathogen infection alters the gene expression landscape of transposable elements in *Drosophila melanogaster*: A meta-analysis

**DOI:** 10.1101/2023.09.03.556151

**Authors:** Sabrina L. Mostoufi, Nadia D. Singh

**Affiliations:** Department of Biology, Institute of Ecology and Evolution, University of Oregon, Eugene, Oregon, United States

**Author notes:** Corresponding author (SLM).

## Abstract

Transposable elements (TEs) are remnants of ancient viral infections and make up substantial proportions of eukaryotic genomes. Current research has begun to highlight the role TEs can play in the immune system response to infections. However, most of our knowledge about TE expression during infection is limited by the specific host and pathogen factors from each study, making it difficult to compare studies and develop broader patterns regarding the role of TEs during infection. Here, we use the tools and resources available in the model, *Drosophila melanogaster*, to analyze multiple RNAseq datasets of flies subject to bacterial, fungal, and viral infections. We analyzed differences in pathogen species, host genotype, host tissue, and sex to understand how these factors impact TE expression during infection. Our results highlight both shared and unique TE expression patterns between pathogens and suggest a larger effect of pathogen factors over host factors for influencing TE expression.

**AUTHOR SUMMARY:** Transposable elements (TEs) are short, repetitive genetic elements that make up substantial proportions of the genomes of all eukaryotic species. These TEs can become expressed during infections, either as a by-product of the infection itself or as a part of the defense mechanism of the immune system. Many studies have investigated TE activity during infection, often focusing on a single pathogen species, one specific host species and genotype, or even a small number of specific TEs. Though these studies provide important detail on these particular cases, less is known about broader patterns of TE activity during infections. Is TE expression the same across different types of infections, and how does the host affect these expression patterns? Here, we analyzed multiple gene expression datasets from infected flies to uncover these broader patterns in TE expression. We found that the type of infection, whether bacterial, fungal, or viral, has a large effect on TE expression. We also found that host factors, like sex, tissue, or genotype, have a much smaller effect on TE expression during infection compared to infection type. Our results offer a broader perspective on how TEs are activated during infections in flies.

## INTRODUCTION

Transposable elements (TEs), or transposons, were first discovered by Barbara McClintock as genetic elements capable of moving around the genome and affecting kernel coloration in *Zea mays* (McClintock 1950). TEs are believed to have originated from ancient viral DNA that became integrated into a host’s genome during a viral infection. TEs make up anywhere from 5-90% of eukaryotic genomes (Guio and González 2019). Though the field originally adhered to the belief that TEs were selfish elements that needed to be transcriptionally silenced, we are now beginning to understand the crucial roles TEs play in genome evolution, gene regulation, and host development for almost all eukaryotes (for review, see Cosby, Chang, and Feschotte 2019).

Today, TEs are organized via several levels of classification. The highest level is class and relates to the method of TE transposition. The most common classes are Class I or DNA transposons, which include TEs that use transposase to invade host genomes, and Class II or RNA transposons, which use reverse transcriptase to replicate throughout a host genome via an RNA intermediate. Within classes, TEs are further classified into families based on sequence similarity and evolutionary history. Finally, at the individual level, different TEs vary in their copy number and locations across host genomes.

There are several potential fates for TEs. In one pathway, TEs accumulate mutations and deletions to eventually become silent. However, there are two other pathways in which TEs can persist: (1) hosts develop ways to suppress disruptive TEs, or (2) TEs mutate and adaptation enables the TE to become an integral part of a host’s gene expression networks (for review, see Cosby *et al*. 2019). TEs that still retain the ability to transpose themselves throughout the genome are generally considered disruptive for their potential to insert themselves into gene regions and interrupt proper gene transposition. To combat the negative effects associated with these disruptive TEs, hosts have developed methods for silencing their expression or preventing their free transposition. One of the most widespread methods is small RNA-mediated silencing of TEs. An example of an sRNA method is the piRNA pathway in *Drosophila melanogaster,* which uses degraded copies of TEs to form piRNA clusters that regulate TE expression in the germline (Kelleher *et al*. 2020). Additionally, ectopic recombination between homologous TEs can result in negative selection against TEs with high copy numbers, long sequences, or which transpose into highly active regions of the genome (Petrov *et al*. 2003; Petrov *et al*. 2011; Kelleher *et al*. 2020). Due to these mechanisms of TE control, many TEs in *D. melanogaster* are found in areas with low recombination and few genes.

However, not all TE insertions are harmful. Several cases have been documented where insertions of TEs disrupt gene functions in surprising ways and create new and beneficial phenotypes (for review, see González and Petrov 2009). In one example, the insertion of the TE *Doc1420* into the gene *CHKov1* created a new, truncated allele that confers pesticide resistance in *D. melanogaster* (Aminetzach, MacPherson, and Petrov 2005). Others have shown that TE insertions into the gene *hsp70Ba* promoter are associated with changes in thermotolerance and female reproductive success in some *D. melanogaster* populations (Michalak *et al*. 2001; Lerman *et al*. 2003; Lerman and Feder 2005). Indeed, there are several TEs which have a high population frequency and display evidence for contributing to adaptation of *D. melanogaster* populations spreading out of Africa (González *et al*. 2008).

Additionally, some TEs have instead evolved to become integrated into host gene networks. In many cases, TEs have taken on the role of promoter, enhancer, or insulator and impact the expression of nearby genes. As an example, in *D. melanogaster* TEs from the *Ty3* (*gypsy*) family can act as promoters or insulators for various genes depending on the genomic location of the TE insertion (Moschetti *et al*. 2020). TEs can also impact gene expression via changes in methylation, as seen in *Arabidopsis thaliana* (Stuart 2016). TEs can also take on larger roles in cellular processes beyond promoters. A well-known example of this phenomenon in *D. melanogaster* are the elements *TART* and *HeT-A*, which are major components of the telomere elongation system in flies (Moschetti 2020).

These systems of regulation and suppression of TEs can become disrupted when the host experiences novel environmental conditions. As early as 1984, McClintock hypothesized that TE activation could occur in response to genome challenges (McClintock 1984). Evidence to support this hypothesis can be found in several systems. For example, in *Caenorhabditis elegans,* heat shock and aging can increase the expression of some TEs (Li *et al*. 2021; Kurhanewicz *et al*. 2020). Temperature stress, including both heat shock and low temperature exposure, impacts TE expression in both *D. simulans* and *D. melanogaster* (Vasilyeva *et al*. 1999; Viera and Biémont 1996; Giraud and Capy 1996; Ratner *et al*. 1992). Additionally, nutrient deficiencies can also lead to TE activation in *Escherichia coli* (Hall 2000) and the wheat pathogen *Zymoseptoria tritici* (Fouché *et al*. 2020).

The effect of infection on TE expression is particularly interesting, given the nature of how TEs became integrated into host genomes initially. Prior to discovering the roles TEs play in host gene networks, many in the field initially theorized that hosts repressed all TEs, but that infection could reawaken those elements. This can be true in some cases, but the hypothesized mechanism is via deregulation, rather than reactivation. For example, in both *Drosophila* and *Rattus* species, the piRNA regulatory pathway can become saturated with pathogenic RNA during a viral infection, resulting in de-repression of native TEs (Durdevic *et al*. 2013; van Gestel *et al*. 2014; Roy *et al*. 2020). However, not every pathogen, host, or TE react the same, and the field has more recently seen TEs play a crucial role in immune system function during pathogen infection. In this case, TEs are upregulated early in viral infection as part of the immune response in both humans and mice (Macchietto *et al*. 2020), as well as in *Drosophila* (Roy *et al*. 2020; Tassetto *et al*. 2017).

Despite the many ways TEs are affected by and can affect the response of their host to infections, efforts to understand these interactions have yet to capture the broader dynamics of the system. Although many studies have investigated the response of TEs to infection, these studies have been generally limited to one host and one infection, and sometimes even a limited number of specific TEs. In particular, studies of TE expression during viral infection are numerous, but investigations using other types of pathogens are scarce. Studies also often analyze different host species, genotypes, sexes, or tissue samples, as well as different pathogen species or strains. These differences make it difficult to understand the broader patterns of TE activation outside of a few, very specific circumstances, even in model organisms. One potential solution to this problem would be to conduct a large-scale study which directly measures the changes in TE expression within a single model organism while altering the type of infection (e.g. viral versus bacterial), host genotypes, and other factors of interest. However, the time, expense, skills, and facilities required for such a study can present significant barriers. An alternative, and more feasible, solution would be to compare numerous studies using the same model organism, allowing us to begin untangling the impact of these host and pathogen factors on TE expression. The fruit fly, *D. melanogaster*, is an ideal candidate for investigating these dynamics because of the number of tools available for TE annotation and the number of gene expression datasets available.

Here, we investigate broad-scale patterns in TE expression during infection of the model organism, *D. melanogaster*. We gathered RNAseq samples from published datasets of *D. melanogaster* infected with a broad range of bacterial, fungal, and viral pathogens. We measured TE expression between control and infected samples and compared patterns both within and between pathogen groups to assess the effect of pathogen species, host genotype, host tissue sample, and host sex on TE expression. Our results show the effect of pathogen on changes in TE expression to be much greater overall than the effect of host factors. We also find that bacterial infections differ significantly from other types of infections, but that infections with the bacterium, *Wolbachia pipientis* look more similar to viral and fungal infections. These findings provide critical insight into how host and pathogen factors can impact TE activity during infection in *D. melanogaster*.

## RESULTS

In this study, we analyzed changes in transposable element (TE) expression between control and infected *Drosophila melanogaster* using RNA-seq data from 14 published datasets (Table 1). Together, these datasets include 31 different fly genotypes and 19 species of individual and multi-species infections of bacterial, fungal, and viral pathogens. Our analyses revealed 231 differentially expressed transposable elements (DETEs) affected by infection in flies. The full list of DETEs is available in S1 Table. We also analyzed patterns of DETEs at the class and family levels across the different pathogen and host factors present in our datasets, which are described in the following sections.

**Table 1:** The RNA-seq datasets used in this study.

| Paper<br>(BioProject) | Pathogen<br>Group | Species/Strain | Control | Infection | Genotypes | Tissue | Sex |
| --- | --- | --- | --- | --- | --- | --- | --- |
| Dobson, *et al.*<br>2016<br>(PRJNA347655) | Bacteria | *Acetobacter pomorum* (DmCS_004) and *A. tropicalis* (DmCS_006) and *Lactobacillus brevis* (DmCS_003) and *L. fructivorans* (DmCS_002) and *L. plantarum* (DmCS_001) | GF | MS | 1 | Whole Body | Male |
| Guo, *et al.* 2014<br>(PRJNA232924) | Bacteria | Conventional microbiome | GF | CV | 1 | Intestine | Female |
| Troha, *et al.* 2018<br>(PRJNA428174) | Bacteria | *Enterococcus faecalis*, *Erwinia carotovora* (Ecc15), *Escherichia coli*, *Micrococcus luteus*, *Providencia rettgeri*, *P. sneebia*, *Pseudomonas entomophila*, *Serratia marcescens* (Db11), *S. marcescens* (Type), *Staphylococcus aureus* | CV | SS | 1 | Whole Body | Male |
| Elya, *et al.* 2018<br>(PRJNA435715) | Fungi | *Entomophthora muscae* | CV | SS | 1 | Brain, Carcass | Female |
| Moskalev, *et al.*<br>2015<br>(PRJNA295562) | Fungi | *Beauveria bassiana* | CV | SS | 1 | Whole Body | Male |
| Paparazzo, *et al.*<br>2015<br>(PRJNA279177) | Fungi | *Beauveria bassiana* | CV | SS | 4 | Whole Body | Male |
| Ramírez-Camejo and Bayman. 2020 (PRJNA377735) | Fungi | *Aspergillus flavus* | CV | SS | 1 | Whole Body | Female |
| Harsh, *et al.* 2020 (PRJNA533975) | Virus | *ZIKV (MR766)* | CV | SS | 1 | Whole body | Female |
| Roy, *et al.* 2020 (PRJNA540249) | Virus | *Sindbis virus (SINV)* | CV | SS | 2 | Carcass, Ovary | Female |
| Lindsey, *et al.* 2021 (PRJNA682591) | Virus, Wolbachia | *Sindbis virus (SINV), Wolbachia pipientis (wMel2)* | CV | SS, MS | 1 | Whole body | Female |
| Detcharoen, *et al.* 2021 (PRJNA602188) | Wolbachia | *Wolbachia pipientis (wMel)* | CV | SS | 1 | Whole body | Female |
| Frantz, *et al.* 2023 () | Wolbachia | *Wolbachia pipientis* | CV | SS | 4 | Ovary | Female |
| Grobler, *et al.* 2018 (PRJNA483452) | Wolbachia | *Wolbachia pipientis* | CV | SS | 1 | Cell culture | Unknown |
| He, *et al.* 2019 (PRJNA439370) | Wolbachia | *Wolbachia pipientis* | CV | SS | 1 | Ovary | Female |

For each dataset, information on the pathogen group (Bacteria, Fungi, Virus, or Wolbachia), the specific species or strain of pathogen, the “control” and “infection” treatments of each study, the number of host genotypes included, and the sample tissue and sex of flies are described. Uninfected flies with a conventional microbiome are represented by “CV”, germ-free flies are represented by “GF”, single-species infections are represented by “SS”, and multi-species infections are represented by “MS.”

Of these 231 total DETEs, 21 were unclassified with an unknown TE class and family. Though these unknown TEs represented approximately 10% of the DETEs in our study, their unclassified status makes it difficult to comment on their influence on host biology and we have excluded them from the rest of our analyses. Future studies may reveal more details about these TEs and their role in host gene networks.

Exclusion of unclassified TEs resulted in a total of 210 DETEs that we used to assess the effect of pathogen and host factors on TE expression. Of all DETEs, 18 came from fungal infection datasets, 47 came from viral infection datasets, and 145 came from bacterial infection datasets, of which 116 were from *Wolbachia pipientis* infections and 29 from non-*Wolbachia* infections (Fig 1). Due to these differences in DETEs among the datasets, we tested whether there was a correlation between sample size and the number of significant TEs within each dataset. We used a linear model to test for this potential relationship and found no significant correlation between dataset sample size and the number of significant TEs (R^2^ = -0.0016, p-value = 0.34).

**Fig 1:**
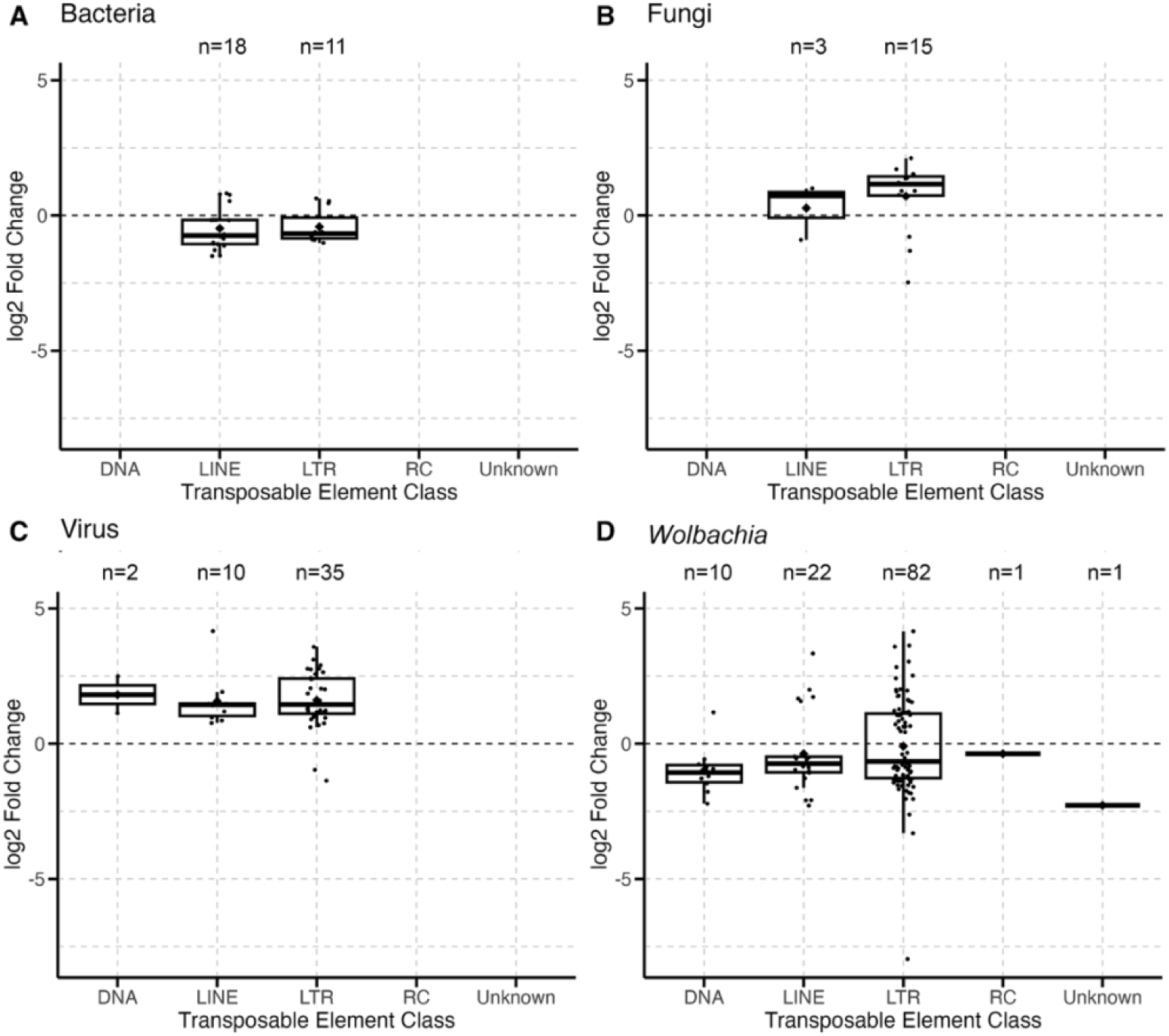
Different infections significantly impact classes of TEs expressed during infection in *D. melanogaster*. Each point represents an individual TE with significant differential expression during infection by either bacteria (A), fungi (B), virus (C), or *Wolbachia* (D). Boxplots present summary statistics, where the top and bottom edges encompass the first to third quartiles and the middle bar represents the median for each group. Boxplot whiskers extend to the smallest and largest nonoutliers. The black diamond in each box represents the average TE expression fold change for each data group.

### Comparisons of Infection by Different Pathogens

To understand how infection affects TE activity, we compared TE expression between flies infected with different pathogens, including bacterial, fungal, and viral infections. We analyzed DETEs at the level of class and family classification and the change in expression, whether increased or decreased, in order to identify larger patterns in TE activity. Due to the large number of datasets with *Wolbachia pipientis*, and because the native strain of *Wolbachia*, *wMel*, is generally not considered pathogenic in *D. melanogaster* (Fry *et al*. 2004), we considered *Wolbachia* infection separately from other bacterial infections.

First, we tested whether there were significant differences in patterns of TE expression between different types of pathogens. Bacterial infections significantly affected different classes and families of TEs compared to all other infection types (Table 2). Fungal, viral, and *Wolbachia* infections affected similar TE classes and families, and there were no significant differences among these groups (Table 2). Pathogen groups also significantly differed in the proportion of TEs with increased or decreased expression during infection (χ-squared = 61.36, df = 3, p-value << 0.00001). Bacterial and *Wolbachia* infections caused 66% of DETEs (n=96) to decrease in expression while fungal and viral infections caused 91% of DETEs (n=59) to increase in expression.

**Table 2:**
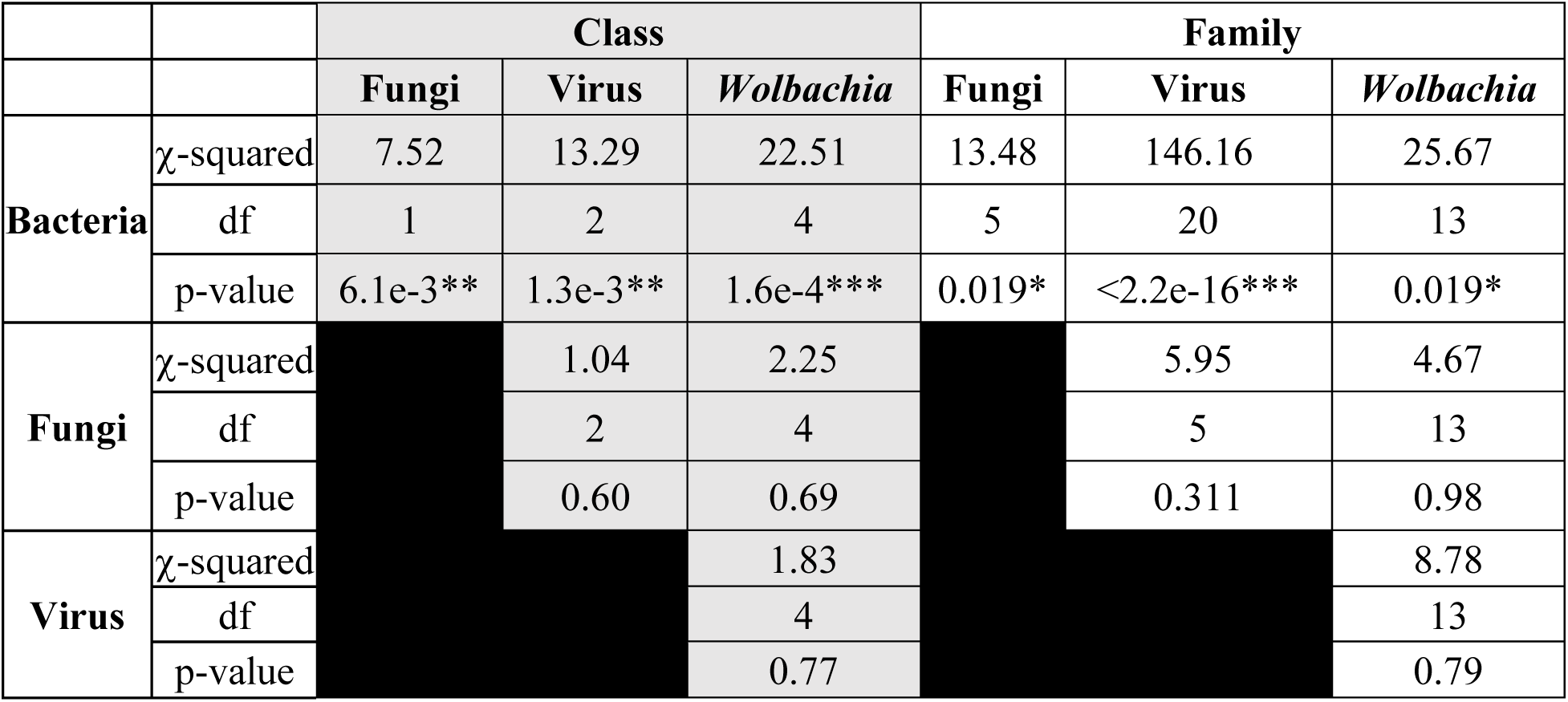
Results from chi-square comparisons of class and family TE proportions between pathogen groups.

|  |  | Class |  |  | Family |  |  |
| --- | --- | --- | --- | --- | --- | --- | --- |
|  |  | Fungi | Virus | Wolbachia | Fungi | Virus | Wolbachia |
| Bacteria | χ-squared | 7.52 | 13.29 | 22.51 | 13.48 | 146.16 | 25.67 |
|  | df | 1 | 2 | 4 | 5 | 20 | 13 |
|  | p-value | 6.1e-3** | 1.3e-3** | 1.6e-4*** | 0.019* | <2.2e-16*** | 0.019* |
| Fungi | χ-squared | | 1.04 | 2.25 | | 5.95 | 4.67 |
|  | df |  | 2 | 4 |  | 5 | 13 |
|  | p-value |  | 0.60 | 0.69 |  | 0.311 | 0.98 |
| Virus | χ-squared | | | 1.83 | | | 8.78 |
|  | df |  |  | 4 |  |  | 13 |
|  | p-value |  |  | 0.77 |  |  | 0.79 |

The proportions of differentially expressed TEs, classified by TE class or family, were compared between each pathogen group. Each group in the table represents the results from a chi-square test between the pathogen groups in the associated row and column for either TE class or TE family proportions. Asterisks indicate significance: * < 0.05, ** < 0.01, *** < 0.001.

Viral, fungal, and *Wolbachia* infections all affected TEs primarily from the LTR class, with *Ty3* being the most abundant TE family represented. However, LTR elements make up approximately 50% of TEs in the *D. melanogaster* genome, 35% of which are from the *Ty3* family. This raised the question of whether these types of infections specifically affected LTR and *Ty3* TEs, or whether their expression was a result of broad and nonspecific activation of TEs during infection. Therefore, we tested whether the class and family proportions of TEs differentially expressed during each infection differed from proportions of TEs present in the *D. melanogaster* genome.

### Effect of Fungal Infections on TE Expression

Our analyses of fungal infections included four datasets of single-species infections, including *Aspergillus flavus*, *Beauvaria bassiana*, and *Entomophthora muscae* (Table 1). Comparison of control and infected samples resulted in 18 DETEs. Fungal infections showed a strong bias for increasing expression of DETEs, where 75% (n=14) of all DETEs increased in expression during infection. Of the 18 DETEs, 83% (n=15) belonged to the LTR class (Fig 1B), and 72% (n=13) came from the *Ty3* family (Fig 2B).

**Fig 2:**
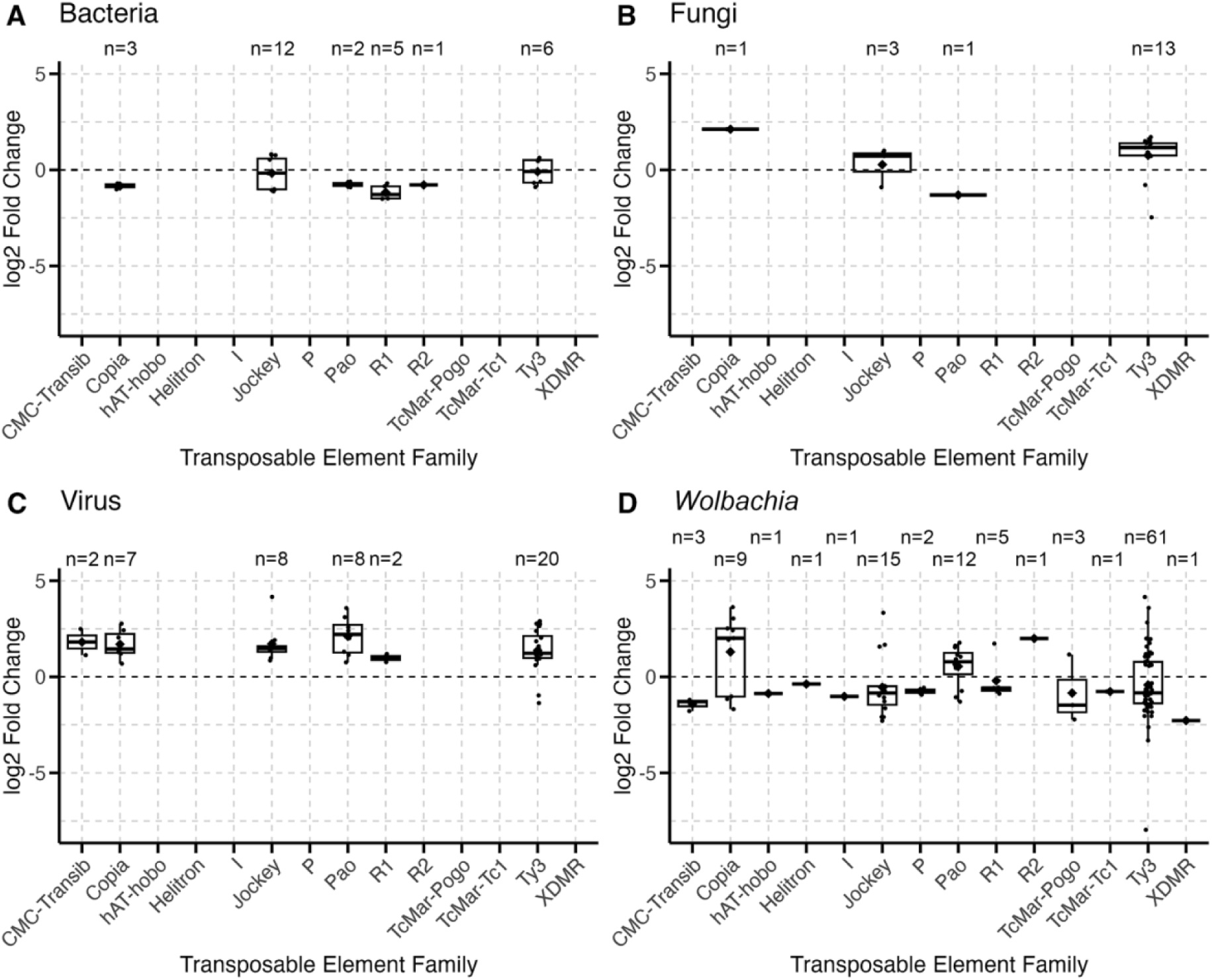
Different infection significantly impact family identity of TEs expressed during infection in *D. melanogaster.* Each point represents a differentially expressed TE during infection by bacteria (A), fungi (B), virus (C), or *Wolbachia* (D). Boxplots present summary statistics, where the top and bottom edges encompass the first to third quartiles and the middle bar represents the median for each group. Boxplot whiskers extend to the smallest and largest nonoutliers. The black diamond in each box represents the average TE expression fold change for each data group.

We also evaluated whether fungal infections broadly affected TEs relative to genomic proportions, or if certain classes or families of TEs specifically respond to fungal infection. To test this, we conducted a chi-square goodness of fit test to compare the proportions of DETE class and family identity with the genomic proportions of TE classes and families in the *D. melanogaster* genome. At the class level, DETEs were not significantly different from the genomic proportions of TE classes (χ-squared = 10.99, df=8, p-value=0.20). This was also true at the family level, where DETEs did not differ significantly from expected family proportions (χ-squared = 15.98, df = 30, p-value = 0.98).

### Effect of Viral Infections on TE Expression

To assess the effect of viral infections on TE expression, we analyzed three RNAseq datasets which included infection with the Zika virus (ZIKV) and Sindbis virus (SINV) (Table 1). Across the datasets, there were 47 DETEs, 96% (n=45) of which increased in expression during infection. The most common class was the LTR class, representing 74% (n=35) of DETEs (Fig 1C), and the most common family was *Ty3*, representing 43% (n=20) of DETEs (Fig 2C).

We also evaluated whether viral infections generally affected TEs, or specifically affected certain TE classes or families. At the level of class, DETEs differed significantly from expected genome proportions (χ-squared = 19.63, df = 8, p-value = 0.01), with residuals indicating that there were more TEs from the LTR class (Chi-square, residual = 2.72) and fewer TEs from the RC class (Chi-square, residual = -2.60) than expected. The family classifications of DETEs also significantly differed from genome proportions (χ-squared = 52.62, df = 30, p-value = 0.0065), with residuals indicating that there were more TEs from the *copia* family (Chi-square, residual = 5.24) and fewer from the *Helitron* family (Chi-square, residual = -2.60) than expected.

### Effect of Bacterial Infections on TE Expression

We analyzed one dataset comparing infection with ten different bacterial species in *D. melanogaster*, and two datasets with conventional and germ-free flies (Table 1). With respect to the former, infection with novel pathogenic and non-pathogenic bacterial species resulted in 28 DETEs, 75% (n=22) of which displayed decreased expression during infection. Over 60% (n=17) of DETEs belonged to the LINE class (Fig 1A), and the most common family was *Jockey*, representing 41% (n=11) of DETEs (Fig 2A). With respect to comparisons between conventional and germ-free flies, there was only one DETE.

The class proportions of the 28 DETEs in the novel bacterial infection datasets differed significantly from expected proportions based on the *D. melanogaster* genome (χ-squared = 32.56, df = 8, p-value = 7.39e-05), with more TEs from the LINE class (Chi-square, residual = 4.80) and fewer TEs from the RC class (Chi-square, residual = -2.01) affected by infection compared to TE class proportions in the genome. Family proportions also differed significantly from expected proportions (χ-squared = 88.36, df = 30, p-value = 1.16e-07), with more TEs from the *copia* (Chi-square, residual = 2.67), *Jockey* (Chi-square, residual = 4.59), *R1* (Chi-square, residual = 4.02), and *R2* families (Chi-square, residual = 5.62) and fewer TEs from the *Helitron* family (Chi-square, residual = -2.01) than expected.

Within the ten bacterial species, there were seven gram-negative and three gram-positive species. We tested whether there were differences within this dataset based on Gram staining. We found no significant differences between gram-negative and gram-positive bacteria species for TE class (χ-squared = 0.78, df = 2, p-value = 0.68) or TE family (χ-squared = 10.71, df = 10, p-value = 0.38). In addition, gram-negative and gram-positive bacteria had nearly identical proportions of TEs with increased or decreased expression, with approximately 75% of TEs decreasing in expression during infection.

### Effect of Wolbachia Infections on TE Expression

We analyzed five datasets of *Wolbachia* infection in *D. melanogaster*, including the *wMel* and *wMel2* variants and one dataset of co-infection between *Wolbachia* and SINV (Table 2.1). Across the datasets of *Wolbachia*-only infections, there were 116 DETEs, 71% (n=82) of which belonged to the LTR class (Fig 1D), and 53% (n=61) of all DETEs were from the *Ty3* family (Fig 2D). Our analyses identified no DETEs in flies co-infected with *Wolbachia* and SINV.

Similar to other bacterial infections, *Wolbachia* infection caused a majority of TEs to decrease in expression, with approximately 64% (n=74) of DETEs decreasing in expression during infection. We found that class proportions of DETEs were significantly different from the proportions of TEs found in the *D. melanogaster* genome (χ-squared = 36.21, df = 8, p-value = 1.61e-05), with more TEs from the LTR class (Chi-square, residual = 3.69) and less from the RC (Chi-square, residual = -3.84) and Satellite classes (Chi-square, residual = -2.37) than expected. Family proportions also differed from expected genome proportions (χ-squared = 98.59, df = 30, p-value = 3.10e-09), specifically that there were more *copia* (Chi-square, residual = 3.45), *R2* (Chi-square, residual = 2.49), *TcMar-Pogo* (Chi-square, residual = 5.90), and *Ty3* TEs (Chi-square, residual = 3.13) and less *CR1* (Chi-square, residual = -2.11), *Helitron* (Chi-square, residual = -3.84), and *Satellite* TEs (Chi-square, residual = -2.37) than expected.

### Comparison of Host Factors during Infection

Across the datasets we analyzed, samples varied by host sex, genotype, and tissue. To assess the effect of these factors on TE expression during infection, we compared the expression of TEs during infection between samples of different host sex, different host genotypes, and different tissue samples. Similar to our analyses of infection, we analyzed DETEs at the level of class and family classification and the direction of changes in expression between different host factors during infection.

### Effect of Host Sex on TE Expression During Infection

First, we analyzed whether host sex had a significant effect on TE expression during infection. Our datasets included ten datasets with samples from female flies, four datasets of male flies, and one dataset of *D. melanogaster* cell culture samples of an unspecified sex. Of the 210 total DETEs analyzed in our study, female samples accounted for 133 DETEs, compared to 30 DETEs in male samples and 47 DETEs in samples of unknown sex (Fig 3).

**Fig 3:**
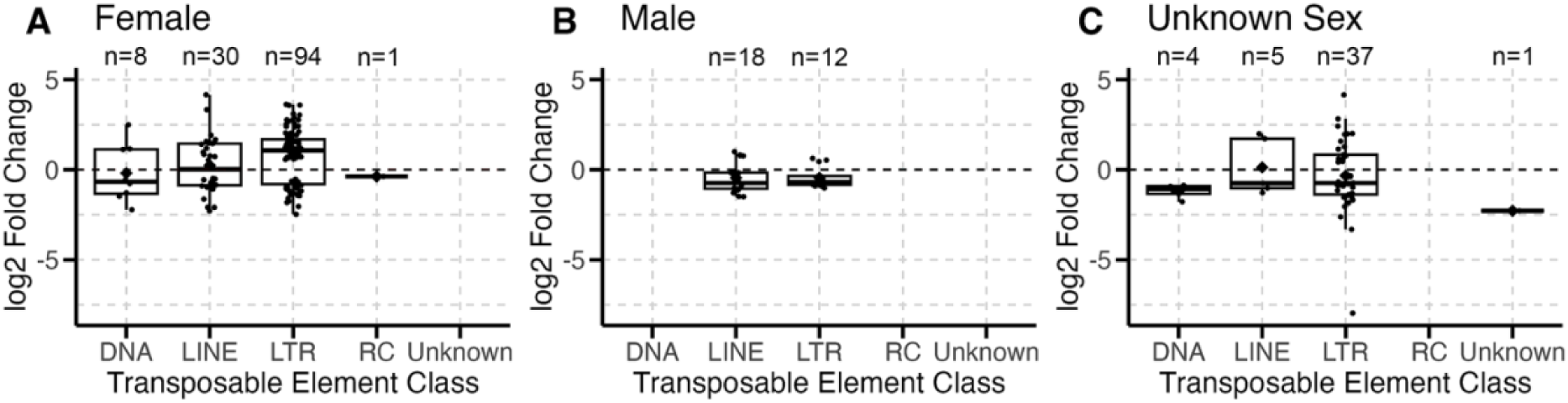
Host sex has a minimal effect on transposable element activity during infection. The number of TEs differentially expressed during infection are grouped by female samples (A), male samples (B), and sample of unknown sex (C). Boxplots present summary statistics, where the top and bottom edges encompass the first to third quartiles and the middle bar represents the median for each group. Boxplot whiskers extend to the smallest and largest nonoutliers. The black diamond in each box represents the average TE expression fold change for each data group.

Over 60% (n=84) of DETEs in female samples showed increased expression during infection, compared to male and unknown sex samples in which 68% (n=53) of DETEs decreased in expression during infection. These differences in expression were significantly different between samples of different host sexes (χ-squared = 21.19, df = 2, p-value = 2.51e-05). Host sex also significantly affected the proportions of both the classes (χ-squared = 29.53, df = 8, p-value = 0.00026) and families (χ-squared = 106.92, df = 28, p-value = 3.72e-11) of TEs that were differentially expressed during infection.

However, unequal associations between pathogen groups and host sex in our datasets may have caused the effect of sex to be confounded by the effect of pathogen. For example, when considering only female samples, which were represented in all pathogen groups, patterns of TE expression were more consistent within pathogen groups than across female samples as a whole. To disentangle this association, we tested for the effect of sex in fungal infections, which had equal numbers of female and male fly datasets. We found no significant effect of sex on the direction of DETE expression (χ-squared = 0.01, df = 1, p-value = 0.92), class identity (χ-squared = 0.11, df = 1, p-value = 0.74) or family identity (χ-squared = 6.19, df = 3, p-value = 0.10). However, female samples still had higher counts of DETEs than male samples, consisting of 16 DETEs versus 2 DETEs in the respective sexes.

### Effect of Host Genotype on TE Expression During Infection

Additionally, the samples in our study included 31 different host genotypes (Table 1). However, due to the sample size of each genotype within the Dobson (2016) dataset, we were unable to include it in this particular analysis, reducing our final analysis to 17 different host genotypes. Of these genotypes, only 9 genotypes had differentially expressed TEs during infection, the most prominent of which are presented in Fig 4. Most genotypes displayed a bias toward decreased expression of DETEs during infection, ranging from 53% to 86% of the total DETEs for each genotype. The exception was the genotype *dcr2^-/-^*, which differed greatly from other genotypes where 98% (n=44) of DETEs increased in expression during infection (Fig 4B).

**Fig 4:**
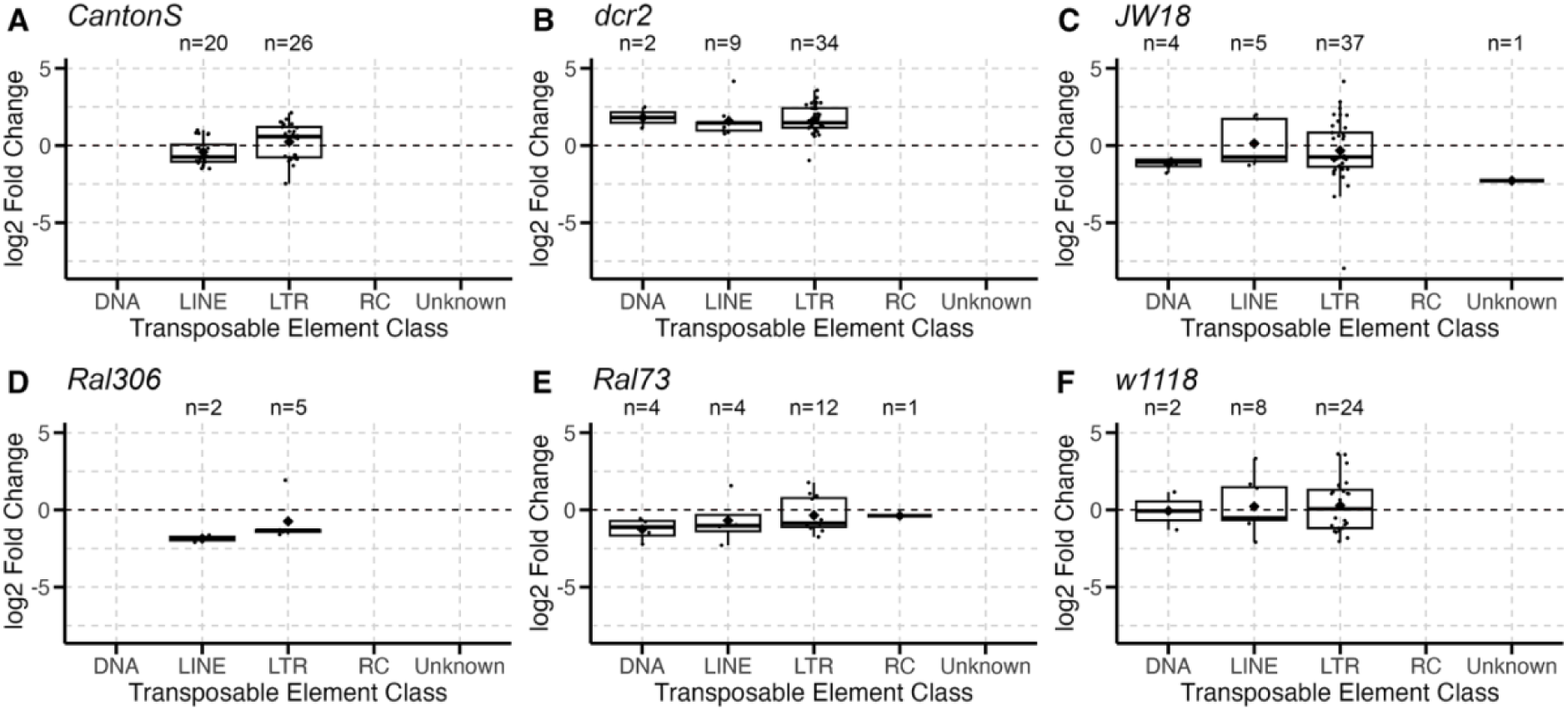
Fly genotype *dcr2* drives effect of host genotype on differences in TE expression during infection. A total of 17 different host genotypes were analyzed for their effect on the classes of TEs expressed during infection in *D. melanogaster*. Only a subset of genotypes with the most differentially expressed TEs are presented in this fig. Each point represents a single differentially expressed TE during infection in the *CantonS* (A), *dcr2* (B), *JW18* (C), *Ral306* (D), *Ral73* (E), or *w1118* (F) fly genotype. Boxplots present summary statistics, where the top and bottom edges encompass the first to third quartiles and the middle bar represents the median for each group. Boxplot whiskers extend to the smallest and largest nonoutliers. The black diamond in each box represents the average TE expression fold change for each data group.

To test for the effect of host genotype on TEs expressed during infection, we used chi square analyses to evaluate differences in expression and whether specific TE classes or families were differentially expressed in different host genotypes. There was a significant effect of host genotype on the direction of TE expression (χ-squared = 59.13, df = 9, p-value = 1.97e-09). However, we found that host genotype did not significantly affect class proportions (χ-squared = 45.54, df = 36, p-value = 0.13) or family proportions of DETEs (χ-squared = 92.71, df = 117, p-value = 0.95) compared to proportions in the *D. melanogaster* genomes.

However, this finding was again potentially related to correlations between host genotype and pathogen group, specifically in the *dcr2^-/-^* genotype, which was only found in one viral infection dataset. When the data were reanalyzed after excluding samples with the *dcr2^-/-^* genotype, host genotype did not significantly impact TE expression (χ-squared = 10.39, df = 8, p-value = 0.24).

### Effect of Tissue Sample on TE Expression During Infection

Finally, we also examined the effect of tissue sample on TE expression during infection. Our datasets included samples from six different tissues, including brain, carcass (whole-body with ovaries removed), cell culture, intestines, ovaries, and whole-body flies (Table 1).

We tested whether DETEs varied significantly by host tissue and found that tissue did not significantly affect the class proportions (χ-squared = 25.36, df = 20, p-value = 0.19) (Fig 2.4C) or family proportions of TEs (χ-squared = 55.27, df = 65, p-value = 0.80). Host tissue significantly affected the direction of expression of DETEs (χ-squared = 51.73, df = 5, p-value = 6.13e-10), specifically that 46 out of 48 DETEs in carcass samples had increased expression during infection (Fig 5B). TE expression was similar across cell culture, ovary, and whole-body samples, with approximately 63% (n=98) of TEs displaying decreased expression during infection (Fig 5).

**Fig 5:**
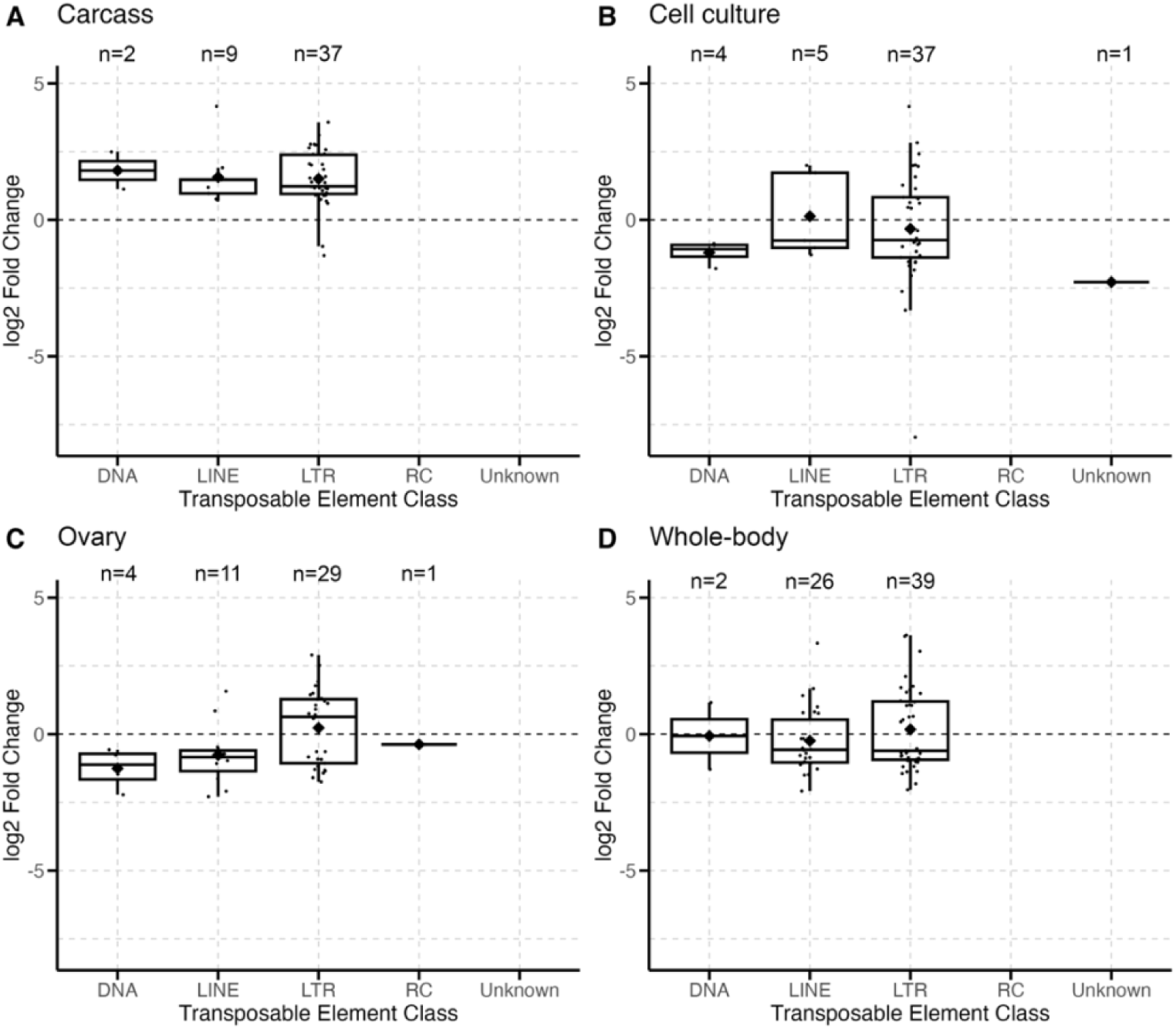
Minimal differences in patterns of TE expression across fly tissue samples during infection. Each point represents a single differentially expressed TE during infection in brain (A), carcass (B), cell culture (C), intestine (D), ovary (E), or whole-body (F) fly tissue samples. Boxplots present summary statistics, where the top and bottom edges encompass the first to third quartiles and the middle bar represents the median for each group. Boxplot whiskers extend to the smallest and largest nonoutliers. The black diamond in each box represents the average TE expression fold change for each data group.

However, similar to our analyses of host sex, there were unequal associations between pathogen groups and host tissue, especially in carcass samples which primarily came from viral infection datasets. Therefore, we reanalyzed the effect of host tissue after excluding carcass samples and found no significant difference in TE expression between tissues (χ-squared = 4.83, df = 4, p-value = 0.31).

## DISCUSSION

In this study, we investigated patterns of transposable element (TE) expression during infection in *D. melanogaster*. Other studies have investigated the effect of infection on TE expression in a variety of hosts and conditions, but experimental differences among these studies make it difficult to directly compare their findings and draw broader conclusions about TE dynamics during infection. By bringing together *D. melanogaster* datasets of differing host genotypes, tissue types, sexes, and pathogens, we sought to uncover the influence of these factors on differentially expressed TEs (DETEs) and identify both unique and broad trends in TE expression during infection.

### Impact of Infection on TE expression

To examine the effect of different infections on TE expression, we examined *D. melanogaster* samples infected with different species of fungal, viral, and bacterial pathogens. Across the different datasets analyzed in this study, infection broadly affected TE expression in flies, though we observed unique differences in the types of TEs and the change in expression depending on the type of pathogen.

First, we analyzed broad trends in whether TEs displayed increased or decreased expression during infection and observed different patterns depending on pathogen group. Samples from viral infections showed a strong bias for increased TE expression, with over 95% of DETEs increasing in expression during infection (Fig 1C). This matches nicely with the well-documented relationship between viruses and TEs, which consistently points to increased TE expression during viral infection (for review, see Hale 2022). There are a couple of mechanisms driving this observed relationship, one of which involves the upregulation of TEs during viral infection as part of the host immune response (Hale 2022). However, increased TE expression can also be a result of viral pathogens disrupting piRNA regulatory pathways, either through oversaturation or by directly manufacturing inhibitors, which can result in de-repression of native TEs (Durdevic *et al*. 2013; van Gestel *et al*. 2014; Roy *et al*. 2020).

Our analyses also identified increased expression of specific TEs known to be active during viral infections. In 2 of the 3 viral datasets we analyzed, we observed increased expression of the *Doc* element, which is known to play a role in virus resistance (Magwire *et al*. 2011; Barrón *et al*. 2014). There were also striking differences between viral strains, where infection by SINV resulted in drastically more DETEs than infection by ZIKV. Both SINV and ZIKV infections are typically non-lethal in *D. melanogaster*, and viral load is controlled by different host immune pathways between the two viral strains (Xu and Cherry 2014; Liu *et al*. 2018). These differences in immune gene response and our results suggest that these viral strains may also differentially influence TE expression in *D. melanogaster*.

Similar to viral infections, we also observed more TEs with increased expression during infection with fungal pathogens (Fig 1B). Fungal infections in the tobacco plant (*Nicotiana tabacum*) can cause increased expression of the *Tnt1* retrotransposon (Grandbastien *et al*. 1997). TE insertions can also play a role in gene duplications that are associated with increased resistance to fungal pathogens (Tan *et al*. 2021), suggesting a potential connection between increased TE expression and the host response to fungal infections. However, there is currently less in the literature about how TEs in *D. melanogaster* respond to fungal infection, making it difficult to draw direct comparisons between the TE response in plants during infection to our findings in flies. Nevertheless, our results suggest that, like in viral infections, fungal infections also lead to increased expression of TEs.

We observed the opposite pattern in bacterial infections, where a majority of TEs displayed decreased expression during infection with various bacterial species, including the endosymbiont, *Wolbachia* (Fig 1). TE insertions are known to affect immune resistance to bacterial pathogens in humans (Bogdan *et al*. 2020) and *D. melanogaster* (Ullastres *et al*. 2021), but it is less clear if TE expression is directly related to bacterial infection resistance. Our results suggest this may be true, based on our comparison between conventional microbiome and novel infection datasets. We analyzed two datasets comparing flies with and without the conventional gut microbiome and found only one significant DETE associated with conventional microbe presence across both datasets. In contrast, novel bacterial infections resulted in several DETEs, suggesting that TE expression is significantly changed during infection and may play a role in the bacterial infection response, as seen in viral infections. Similar to Troha and colleagues (2018) who observed that bacterial pathogenicity did not directly correlate with the number of host genes regulated during infection, we also found that the number of DETEs did not correlate with the severity of bacterial infection.

Following this trend, we observed that *Wolbachia* infections resulted in many DETEs, despite the fact that *Wolbachia* infections are native and non-pathogenic in *D. melanogaster.* Infection with *Wolbachia* was also similar to other bacterial infections by generally decreasing TE expression during infection (Fig 1D). These results agree with others that have found *Wolbachia* infection to decrease TE activity (Touret *et al*. 2014). Another study also showed that *Wolbachia* can differentially affect expression of some TEs in *D. melanogaster*, both increasing and decreasing TE expression depending on TE identity and host genotype (Eugénio *et al*. 2022). While it is plausible that changes in TE expression during infection with other pathogens may be related to host infection responses, this reasoning does not explain why infection with *Wolbachia* should alter TE expression in its host. We discuss more on the effect of *Wolbachia* on its host in the following sections.

These similarities and differences between pathogens, particularly that viral and fungal infections increased TE expression while bacterial infections decreased TE expression, raise interesting questions about whether these patterns are driven by the mechanism of infection for each pathogen. One mechanism to consider is whether each pathogen operates via intracellular or extracellular infections in flies, which are combatted by different immune reactions in *Drosophila* (Kurata 2010). Viruses and most fungi are intracellular pathogens in flies, which could be related to the increase TE expression across both types of infections, while many bacterial infections are extracellular. If TEs are responding as part of the immune system during infection, this could explain why we observed different patterns in TE expression if different immune response networks are activated during different types of infections.

Unfortunately, this correlation breaks down in several ways. First, though *Wolbachia* is an intracellular infection, infections with *Wolbachia* differed from other intracellular fungal and viral infections by decreasing expression of a majority of TEs. *Drosophila* immune responses to fungal pathogens also share more similarities to bacterial pathogens (Lemaitre *et al*. 1997), though we observed opposite patterns in TE expression between fungal and bacterial infections. These interesting patterns in TE expression suggest some divide between bacterial pathogens versus viral and fungal pathogens, though it may not be related to the mode of infection.

### Impact of Infection on TE Class and Family

In addition to the direction of expression, we also analyzed what types of TEs were differentially expressed during infection. Across all infections, most DETEs were RNA retrotransposons, with a small number of DNA transposons differentially expressed in viral and *Wolbachia* infection datasets (Fig 1). Other studies have found that DNA transposons are the most active TEs in *Drosophila simulans* (Kofler *et al*. 2015), but our results agree with others that have found retrotransposons, specifically LTR elements, to be the most active elements in *D. melanogaster* (Kofler *et al*. 2015).

We also observed little or no Rolling Circle (RC) DNA transposons across all infections, and this was significantly less than expected in viral, bacterial, and *Wolbachia* infections. RC elements make up approximately 14% of TEs in the *D. melanogaster* genome, with the RC family *DINE-1* estimated to have the highest copy number of repeats of any TE family (Thomas *et al*. 2014). *DINE-1* elements are often involved in gene duplications, some of which have been linked to insecticide resistance in *Drosophila* (Carareto *et al*. 2014). Yet, despite their prevalence, we observed very few RC elements that were differentially expressed during infection in *D. melanogaster*.

We found that the classes and families of TEs expressed during fungal infection did not significantly differ from genome proportions, suggesting no preference for the types of TEs affected by fungal infections. However, both viral and bacterial infections, including *Wolbachia* infections, affected significantly more TEs from the *copia* family than expected based on the prevalence of these TEs in the genome. *Copia* elements are known to regulate numerous host genes in *Drosophila* related to development (for review, see Moschetti *et al*. 2020). We observed relatively strong expression (> 2 log2-fold change) of *copia* elements across multiple host genotypes and tissue types in both *Wolbachia* and viral infection datasets (S1 Table), suggesting a potential role for these elements during the host response to infection.

Bacterial infections differed significantly from other types of infections and were more likely to affect LINE elements (Fig 1A) and TEs from the *Jockey* family (Fig 2A). Of particular note were the elements *HeT-A* and *TART*, which decreased in expression during bacterial infection across several datasets. Both of these TEs are known to affect telomere elongation and chromosome stability in flies (Capkova Frydrychova *et al*. 2008). Other studies have shown that flies modulate gene expression related to stress and cell homeostasis during bacterial infection (Troha *et al*. 2018), which would include processes relating to telomeres and chromosome structure. Other notable TEs that were differentially expressed during bacterial infection include *invader*, *BURDOCK*, and *BS* elements, which have been linked to expression of immune-related genes that increase infection resistance in *D. melanogaster* (Ullastres *et al*. 2021). Overall, these patterns suggest a decrease in TE activity related to normal cell maintenance processes, perhaps in favor of increasing resources dedicated toward infection resistance.

*Wolbachia* infections differed from infections with other bacterial species, affecting more TEs from the LTR class and the *Ty3* family (Fig 2D). Other studies have found that *Wolbachia* infection can decrease expression of *Ty3* elements in *D. melanogaster* (Touret *et al*. 2014), but this effect may differ depending on host genotype (Eugénio *et al*. 2022). Though we did not find a significant effect of genotype, we observed that the effect of *Wolbachia* infection can differ across TEs and datasets, with *Wolbachia* infection increasing and decreasing the expression of various *Ty3* elements. This finding may relate to *Wolbachia*’s ability to alter host gene expression that has been observed in other studies (He *et al*. 2019; Frantz *et al*. 2023). TEs from the *Ty3* family have been shown to act as promotors and insulators of host genes (Moschetti *et al*. 2020), suggesting a potential connection between *Wolbachia*’s effect on TE expression also resulting in changes to gene expression in its hosts.

*Wolbachia* infection included the most DNA DETEs of any infection type and affected significantly more TEs from the *Tc1/mariner* superfamily than expected. This was of particular interest to us because of *Wolbachia*-associated plastic recombination in *D. melanogaster* (Singh 2019; Bryant and Newton 2020; Mostoufi and Singh 2022). DNA transposons are associated with increased recombination rate in the wood white butterfly (*Leptidea sinapis*) (Palahí I Torres *et al*. 2023) and in *C. elegans* (Duret *et al*. 2000). Additionally, heat shock can cause increased gene expression of the *Tc1-mariner* retrotransposon in *C. elegans*, leading to increased DNA double-strand breaks (Kurhanewicz *et al*. 2020). The mechanism behind how *Wolbachia* alters recombination rate in *D. melanogaster* is currently unknown, but these results suggest a potential connection between increased expression of DNA transposons and increased recombination rate that is facilitated by *Wolbachia*. Future experiments which directly test this hypothesis may shed further light on this potential mechanism for *Wolbachia*-associated plastic recombination.

### Impact of Host Factors on TE Expression During Infection

We also examined how host factors affected TE expression. We analyzed differences between female and male fly samples, in addition to a cell culture of unknown sex, to determine whether host sex significantly changed TE expression during infection. Sex is known to affect the susceptibility and intensity of infections in humans, with women generally less susceptible to infection due to more robust immune responses compared to men (Klein and Flanagan 2016). *Drosophila* males and females also exhibit several differences in their responses to infection, including sex chromosome-linked variation in immune responses (Taylor and Kimbrell 2007; Hill-Burns and Clark 2009), as well as differences in behavioral responses to infection (for review, see Belmonte *et al*. 2020). However, unlike in humans, there is no one sex in *D. melanogaster* which is overall more responsive to infection, but different types of infections are biased towards male or female survival (Belmonte *et al*. 2020). Additionally, TEs are known to be involved in sexual development and other sex-specific forms of gene expression (for review, see Dechaud *et al*. 2019).

When analyzing for the effect of host sex, we found that differences in TE expression were largely driven by pathogen due to sex biases in several of the datasets. Within fungal infections, which had equal proportions of male and female samples, sex did not significantly impact expression or class proportions of DETEs. However, we still observed that female samples had more DETEs than male samples. These findings suggest that host sex may influence the number of TEs differentially expressed during infection, but not necessarily the types of TEs affected or whether expression is increased or decreased. This finding is particularly interesting in light of other reports which demonstrate that males are more likely to survive fungal infections than females (Taylor and Kimbrell 2007; Shahrestani *et al*. 2018). So while our results suggest that female flies may see greater changes in TE expression during infection, these changes may not directly correspond to infection resistance to fungal pathogens. The connection between host sex and TE expression and infection resistance may instead play a larger role in other types of infections, particularly with bacterial or viral pathogens. Further investigations of the effect of sex and TE expression during infections with bacterial or viral pathogens would help to clarify this relationship.

In addition to host sex, we also evaluated the effect of host genotype during infection (Fig 4). We analyzed a total of 17 genotypes and found no significant differences in TE class proportions across host genotypes. Similar to our analysis of sex, host genotype was significantly associated with differences in TE expression, although this was largely driven by one genotype, *dcr2-/-*, found in only viral infection datasets. This large effect is likely due to a mutation in the genotype that impairs siRNA formation and leads to higher viral replication (Roy *et al*. 2020). Another host genotype, *CantonS*, was present in both bacterial and fungal infection datasets, but showed very few similarities within the genotype and was more similar to other fly genotypes within the same pathogen infection group. These results suggest that the influence of non-mutant host genotypes on TE expression is smaller than the influence of the pathogen.

Host genotype is known to influence TE copy number and insertion location between species and populations (Barrón *et al*. 2014; Signor 2020). Therefore, it was expected that TE copy number and location would vary between the genotypes in our datasets. Indeed, we observed differences in base mean counts of the same TEs across different samples. TE insertions and copy number can affect TE expression (Lee and Langley 2012), but the piRNA pathway in *Drosophila* also employs copy-dependent silencing of TEs (Kelleher *et al*. 2020). By directly comparing uninfected and infected flies from the same dataset, our analyses assessed the relative TE expression within each genotype and found that host genotype does not significantly affect the change in TE expression during infection.

Sample tissue also varied across the datasets used in this study, including samples from brain, cell culture, carcass, intestine, ovary, and whole-body samples (Fig 5). Host tissue was associated with significant differences in the direction of TE expression, but differences in TE class and family were nonsignificant. Specifically, carcass tissue samples generally had increased TE expression during different types of infection, while patterns in whole-body and ovary tissues shared more similarity by pathogen group. Outside of infection, hosts differentially regulate TE expression in different tissues, where somatic expression is generally repressed compared to germline expression (Haig 2016). However, one experiment found the opposite to be true during infection. In Roy *et al*. (2020), TE expression during viral infection was much higher in somatic tissues than in germline tissues, which was confirmed during our analysis of the same data.

Taken together, these results suggest a small role of host factors in influencing TE expression during infection. Although host sex and some tissue samples showed shared patterns in the direction of TE expression, other samples showed little consistency within groups and were more similar to samples from the same pathogen group. In contrast, samples from the same type of infection shared many more patterns in TE expression, suggesting that infection type had a stronger influence on TE expression than host factors. These findings differ from other studies analyzing non-TE gene expression during infection, where host genotype had a larger effect than infection status in *D. melanogaster* (Frantz *et al*. 2023) and the type of pathogen infection in humans (Idaghdour *et al*. 2012). These somewhat contradictory results may relate to differences in the ways that host genes and transposable elements are activated or suppressed during infection. A majority of TEs are repressed by host mechanisms, which may become overwhelmed during infection and allow for previously silenced TEs to become active. This has been observed during infections in mammals (van Gestel *et al*. 2014) and flies (Roy *et al*. 2020). Therefore, the type of infection may impact which host defense mechanisms are activated and become unable to regulate TE expression.

## CONCLUSIONS

Transposable elements can play crucial roles in the host’s immune system response to infection, as has been demonstrated in humans, mice, flies, and more. However, our understanding of these interactions is limited by differences between pathogen and host factors in each study, affecting our ability to identify broader patterns in TE responses to infection. Our work presented here combines gene expression data from multiple infection studies in *D. melanogaster* to illuminate the influence of pathogen and host factors on TE activity during infection. We find that TE activity is strongly affected by differences in pathogen infection, while the effect of host factors is comparatively smaller. Future experimental work in flies, as well as additional comparative studies in other model organisms, would help to expand our understanding of TE activity during infection to even broader scales.

## MATERIALS AND METHODS

### Sequence processing

The datasets used in this study were downloaded from the European Nucleotide Archive (ENA) at EMBL-EBI. Processing and analysis of RNA-seq files was completed using the University of Oregon’s high performance computing cluster, Talapas. Raw fastq files were merged and aligned to the *Drosophila melanogaster* reference genome (Release 6.41) as unsorted BAM files using the program STAR (v2.7.9a) (Dobin *et al*. 2013). To optimize the data for transposable element (TE) analysis, we used the recommended settings for STAR from Jin and Hammell (2018) by setting –outFilterMultimapNmax 100 and –winAnchorMultimapNmax 200, with default values for all other parameters.

### Measuring transposable element expression

We used the program TEtranscripts to count TEs in our selected datasets (Jin *et al*. 2014). TEtranscripts requires two GTF files, one for gene sequences and one for TE sequences. We used the *D. melanogaster* TE GTF file provided on the Hammell lab website (labshare.cshl.edu/shares/mhammelllab/www-data/TEtranscripts/TE_GTF/dm6_BDGP_rmsk_TE.gtf) and the *D. melanogaster* genome Release 6.32 GTF file from Ensembl (ftp.ensembl.org/pub/release-107/gtf/drosophila_melanogaster/Drosophila_melanogaster.BDGP6.32.104.gtf.gz). After alignment, BAM files for control and infection groups from each dataset were analyzed using TEtranscripts, with –norm TC and default parameters, to produce TE gene counts.

When presenting our findings, we have also elected not to use the antiquated and problematic name for the *gypsy* TE family, instead opting for the alternate naming of *Ty3* as advocated for by others in the field (Wei *et al*. 2022).

### Statistical analyses

All statistical analyses and data visualization were completed in RStudio (2023.06.0, “Mountain Hydrangea” Release). Differentially expressed transposable elements (DETEs) were identified by calculating the log2 fold change between control and infection samples and assessing significance using DESeq2 (v1.32.0) (Love, Huber, and Anders 2014). We used a chi-squared goodness of fit test to examine whether observed patterns of differentially expressed TEs at class and family levels were significantly different from proportions of TEs present in the *D. melanogaster* genome, as well as to compare differences between and within pathogen groups. Genomic proportions of TEs categorized by class and family were calculated from the *D. melanogaster* genome Release 6.32 Ensembl GTF annotation file.

## ACKNOWLEDGEMENTS

The authors would like to thank Brendan Bohannon, Bill Cresko, Karen Guillemin, and Ken Prehoda for providing comments that improved the quality of analyses in this study. We would also like to thank members of the Singh lab group, Annette Estevez and LyAndra Lujan, as well as Justin Blumenstiel and Erin Kelleher for valuable feedback on the manuscript. Additionally, this work benefited from access to the University of Oregon high performance computing cluster, Talapas.

## SUPPORTING INFORMATION CAPTIONS

**S1 Table. Full list of differentially expressed transposable elements from each dataset.**

